# The PK/PD Integration and Resistance of Tilmicosin Against *Mycoplasma hyopneumoniae*

**DOI:** 10.1101/2020.03.04.977488

**Authors:** Zilong Huang, Zixuan Hu, Xirui Xia, Haorui Zheng, Xiaoyan Gu, Xiangguang Shen, Hong Yang, Huanzhong Ding

## Abstract

*Mycoplasma hyopneumoniae* is the major pathogen causing enzootic pneumonia in pigs. *M. hyopneumoniae* infection can lead to considerable economic losses in the pig-breeding industry. Here, this study established a first-order absorption, one-compartment model to study the relationship between the pharmacokinetics/pharmacodynamics (PK/PD) index of tilmicosin against *M. hyopneumoniae in vitro*. We simulated the drug concentration of timicosin in the fluid lining the lung epithelia of pigs. The minimum inhibitory concentration (MIC) of tilmicosin against *M. hyopneumoniae* with an inoculum of 10^6^ CFU/mL was 1.6 μg/mL using the microdilution method. Static time–kill curves showed that, if the drug concentration >1 MIC, the antibacterial effect showed different degrees of inhibition. At 32 MIC, the amount of bacteria decreased by 3.16 log_10_ CFU/mL, thereby achieving a mycoplasmacidal effect. The *M. hyopneumoniae* count was reduced from 3.61 to 5.11 log_10_ CFU/mL upon incubation for 96 h in a dynamic model with a dose of 40–200 mg, thereby achieving mycoplasmacidal activity. The peak concentration by MIC (C_max_/MIC) and the area under the concentration-time curve over 96 h divided by the MIC (AUC_0–96 h_/MIC) were the best-fit PK/PD parameters for predicting the antibacterial activity of tilmicosin against *M. hyopneumoniae* (R^2^ = 0.99), suggesting that tilmicosin had concentration-dependent activity. The estimated value for C_max_/MIC and AUC_0–96 h_/MIC for 2log_10_ (CFU/mL) reduction and 3log_10_ (CFU/mL) reduction from baseline was 1.44 and 1.91, and 70.55 h and 96.72 h, respectively. Four *M. hyopneumoniae* strains (M1–M4) with reduced sensitivity to tilmicosin were isolated from the four dose groups. The susceptibility of these strains to tylosin, erythromycin and lincomycin was also reduced significantly. For sequencing analyses of 23S rRNA, an acquired A2058G transition in region V was found only in resistant *M. hyopneumoniae* strains (M3, M4). In conclusion, in an *in vitro* model, the effect of tilmicosin against *M. hyopneumoniae* was concentration-dependent and had a therapeutic effect. These results will help to design the optimal dosing regimen for tilmicosin in *M. hyopneumoniae* infection, and minimize the emergence of resistant bacteria.

## 1 Introduction

*Mycoplasma hyopneumoniae* is the primary pathogen of mycoplasmal pneumonia in pigs. *M. hyopneumoniae* is widespread in various regions, and can cause huge economic losses to the pig industry [1]. Infected pigs are the main source of infection. The pathogen can be transmitted directly through air and contact, so the infection rate is extremely high [2]. Once flocks of pigs are infected with *M. hyopneumoniae*, eradication is difficult because *M. hyopneumoniae* can transmit vertically [3, 4].

The first-line method of controlling *M. hyopneumoniae* infection is antibiotic treatment [5]. Tilmicosin is a macrolide antibiotic used commonly in animals. It has a long elimination half-life and high concentration in lung tissues [6, 7]. Tilmicosin has a broad spectrum of efficacy, especially against *Mycoplasma* species [8]. The unique antibacterial mechanism of tilmicosin is perfect for treatment of *M. hyopneumoniae* infections. However, the unreasonable use and abuse of macrolides has led to the emergence of drug resistance [9, 10]. Most of the resistance mechanisms of macrolides include active efflux mechanisms, changes in target molecules bound by drugs, and inhibition of enzyme activities [11]. The resistance to macrolides is associated with mutations in domains II or V of 23S rRNA genes, or the rplD and rplV genes encoding ribosomal proteins L4 and L22 [10, 12, 13].

Due to the difficulty in culturing and counting of *M. hyopneumoniae*, the pharmacokinetic/pharmacodynamic (PK/PD) profiles of tilmicosin against *M. hyopneumoniae* are very limited. Also, establishing an infection model of *M. hyopneumoniae in vivo* is challenging. Therefore, it is a feasible to establish an *in vitro* dynamic model to evaluate the effect of tilmicosin against *M. hyopneumoniae. In vitro* PK/PD models have been used widely to optimize dose regimens, monitor antimicrobial activity, and prevent the emergence of resistant bacteria [14]. Such models can simulate the change in drug concentration in animals but also eliminate differences among animals [15]. Moreover, the PK/PD parameters in the *in vitro* model are very similar to those in the animal-infection model [16-18]. Therefore, establishment of an *in vitro* dynamic model appears to be a viable way to evaluate the effects of tilmicosin on *M. hyopneumoniae*.

We wished to apply the one-compartment infection model *in vitro* to determine the PK/PD indices of tilmicosin against *M. hyopneumoniae*. In this way, we could investigate the mechanism of resistance. This model can be used as a reference to optimize the dosing regimen for tilmicosin against *M. hyopneumoniae*.

## 2 Materials and Methods

### 2.1 Materials

A standard strain of *M. hyopneumoniae* (ATCC 25934) was obtained from the Chinese Veterinary Microorganism Culture Collection Center (Beijing, China) and stored at −80 °C. Tilmicosin (75.8%), tylosin (82.6%), erythromycin (85.0%), tiamulin (99.0%), doxycycline (85.8%), and enrofloxacin (99.0%) were kindly supplied by Guangdong Dahuanong Animal Health Products (Xincheng, China). Amikacin (99.0%) and lincomycin (84.6%) were purchased from Guangdong Puboxing Animal Health Products (Guangzhou, China) and stored at −80 °C before use.

A fresh stock solution (1280 mg/L) of each antibacterial agent was prepared for each experiment. Broth medium base was purchased from Qingdao Hope Biological Technology (Qingdao, China). The reduced form of nicotinamide adenine dinucleotide was obtained from Beijing Newprobe Biotechnology (Beijing, China). L-Cysteine was purchased from Beijing Solarbio Science and Technology (Beijing, China).

### 2.2 Determination of the minimum inhibitory concentration (MIC)

The MIC of tilmicosin against *M. hyopneumoniae* was determined by a modified version of the MIC assay, as described by Tanner and Wu [19]. Briefly, dilutions of exponential-phase cells at 10^5^, 10^6^ and 10^7^ CFU/mL (100 μL) were added to an equal volume of drug-containing medium per well in a 96-well plate. The tilmicosin concentration in the 96-well plate was 0.05∼12.8 μg/mL. Growth control (lacking antibiotic), sterility control (sterile broth at pH 7.7) and endpoint control (blank medium at pH 6.5) were included. Plates were incubated at 37 °C in a humidified atmosphere of 5% CO_2_ until the growth group and endpoint control were the same color. The MIC was defined as the minimal concentration of antibacterial agent that resulted in no color change. All experiments were carried out in triplicate.

According to the method described by Hannan et al. [20], plates containing a series of tilmicosin concentrations (1.6∼25.6 μg/mL) were prepared. Samples (10 μL) of cultures with an inoculum of 10^5^, 10^6^ and 10^7^ CFU/mL were also applied to the drug plates. A blank-growth control group was also used and comprised cells spread on plates lacking the drug. Plates were incubated for ≥7 days. The lowest concentration without *M. hyopneumoniae* growth was determined as the MIC. All experiments were carried out in triplicate.

### 2.3 Time–kill curves

Four milliliters of blank medium, 0.5 mL of 10-times the final drug concentration, and 0.5 mL of logarithmic *M. hyopneumoniae* were added to a bottle in turn and then mixed. The tilmicosin concentration in the culture system was in a certain range (1/2, 1, 2, 4, 8, 16, and 32 the MIC that was determined for an *M. hyopneumoniae* inoculum of 10^6^ CFU/mL). A growth control (not exposed to the drug) and a sterility control (medium at pH 7.7 without the drug or *M. hyopneumoniae*) were indispensable. Penicillin bottles were cultured for 60 h at the environments stated above. Aliquots (100 μL) of the culture were taken from each bottle at 0, 1, 3, 6, 9, 12, 24, 36, 48 and 60 h to detect the *M. hyopneumoniae* population. After 7 days, the results were read using an inverted microscope (Leica, Germany).

### 2.4 PK/PD model *in vitro* and dosing regimens

This study used an *in vitro* dynamic model described previously [18]. This experiment was done to simulate the timicosin concentration in the fluid lining the lung epithelia of pigs [7]. The model was applied according to the outline shown in **Figure 1**.

**Figure 1.**
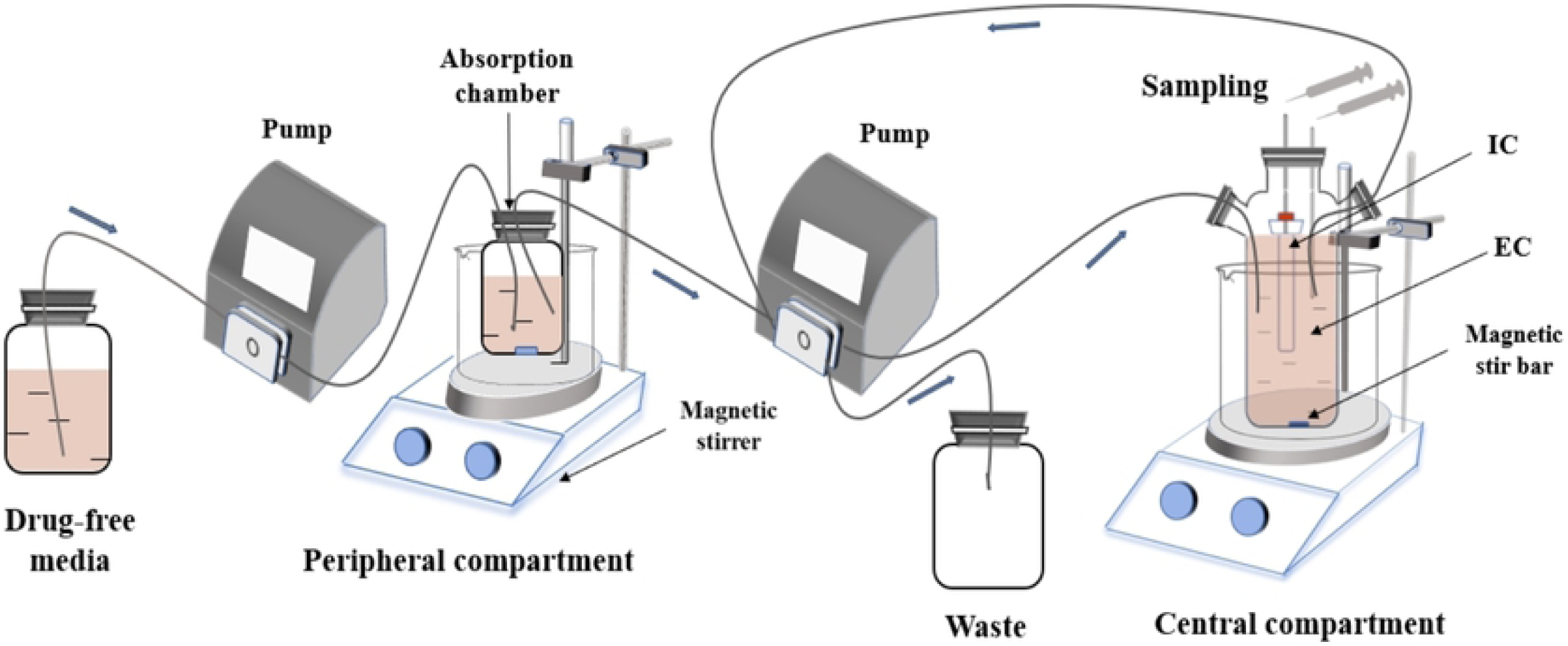
An *in vitro* model that simulates the pharmacokinetics of tilmicosin in the fluid lining the lung epithelia of pigs and determines the effects of tilmicosin on the growth and susceptibility of *M. hyopneumoniae*. EC, external compartment; IC, internal compartment.

Briefly, the model system consists of three parts. The first part is an absorption chamber containing a drug medium as a site of administration. The second part is a central chamber, which comprised 300 mL of sterile medium (external compartment [EC]) and a 10-mL dialysis tube (internal compartment [IC]). The third part is a reserve room for fresh media. At the same time, the waste liquid is collected in the waste liquid chamber.

The model parameters were determined by the colonization site of *M. hyopneumoniae* and PK characteristics of tilmicosin in pigs. The parameter values of absorption half-life, elimination half-life, and flow rate of peristaltic pumps were 12.17 h, 17.16 h and 0.29 mL/min, respectively. According to the clinically recommended dose, eight dose groups (10, 20, 40, 60, 80, 120, 160 and 200 mg) were designed for the *in vitro* dynamic model. Increasing the speed of the magnetic stirrer as the drug was injected into the absorption chamber achieved a rapid balance between the inside and outside of the dialysis membrane simultaneously.

Samples (2 mL) were collected from the EC 1, 3, 6, 9, 12, 24, 36, 48, 72 and 96 h after administration, and then stored at −20 °C until analyses. Samples (100 μL) were taken from the IC before dosing as well as 6, 12, 24, 36, 48, 72 and 96 h after administration. Collected samples were used to detect the number of, and susceptibility to, *M. hyopneumoniae*.

### 2.5 Determination of the tilmicosin concentration in the medium

The tilmicosin concentration in the medium was analyzed using a high-performance liquid chromatography unit (1200 series; Agilent Technologies, Santa Clara, CA, USA) and a triple quadrupole mass spectrometer (6410; Agilent Technologies) equipped with an electrospray ionization source. The analytical method used was as described by Huang et al. [18]. The standard curve (R^2^ > 0.99) was defined by six calibration standards of tilmicosin with a final concentration ranging from 5 ng/mL to 500 ng/mL. The limit of detection (LoD) and limit of quantification (LoQ) in the medium was 0.5 ng/mL and 1 ng/mL, respectively.

### 2.6 Integration and modeling of PK/PD

Three important PK/PD indices were calculated by integrating PK parameters and MIC value (MIC=1.6 μg/mL) *in vitro*: the peak concentration by MIC (C_max_/MIC), the area under the concentration–time curve over 96 h divided by the MIC (AUC_0–96 h_/MIC) and the cumulative time that the concentration exceeds the MIC (%T > MIC).

The correlation between PK/PD indices and antimicrobial activity against *M. hyopneumoniae* was analyzed using WinNonlin (Certara, Princeton, NJ, USA). We chose the inhibitory sigmoid *E*_max_ model to analyze data:

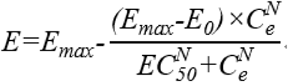

where *E* is the anti-*M. hyopneumoniae* effect; *E*_max_ is the change in the amount of *M. hyopneumoniae* in the control group at a 96-h interval; *E*_0_ is the largest anti-*M. hyopneumoniae* effect, determined as log_10_CFU/mL reduction at the same interval; *C*_*e*_ represents the PK/PD indices (%T > MIC, C_max_/MIC and AUC_0–96 h_/MIC); *N* is the Hill coefficient that describes the steepness of the PK/PD indices–effect curve; *EC*_*50*_ is the corresponding PK/PD value when the anti-*M. hyopneumoniae* effect reaches 50% of the maximum antibacterial effect; *R*^*2*^ was calculated for each assay.

### 2.7 Susceptibility testing of *M. hyopneumoniae* and DNA sequencing

This study used the method described by Huang et al. [18] with slight modification. Briefly, 100 μL of the bacteria at the final time point in the IC qas collected. Then, every 10 μL of the bacterial solution was placed on the surface of the drug plate containing 1× MIC concentration. After 7 days of culture, colonies that resumed growth were transferred to blank liquid medium and sub-cultured five times until their growth was stable. The MIC of these strains was re-determined, and colonies with reduced sensitivity to tilmicosin were screened. After five generations, amplification by polymerase chain reaction and sequencing of stabilized MIC mutants was carried out using Sanger sequencing by TsingKe Biological Technology (Chengdu, China). The sensitivity of these strains to other antimicrobial agents (lincomycin, amikacin, enrofloxacin, doxycycline, tiamulin, erythromycin, and tylosin) was also tested.

## 3 Results

### 3.1 Susceptibility determination

The MIC of tilmicosin against *M. hyopneumoniae* with inoculums of 10^5^, 10^6^, and 10^7^ CFU/mL was 0.8, 1.6, and 1.6 μg/mL using the microdilution method, and 3.2, 6.4, and 6.4 μg/mL using the agar dilution method, respectively.

### 3.2 Analyses of time–kill curves

The *in vitro* static bactericidal curves of different concentrations of tilmicosin against *M. hyopneumoniae* are shown in **Figure 2**. When the drug concentration was 0.5 MIC, the bacteria continued to grow, and the number of bacteria increased by 1.31 log_10_ CFU/mL. The number of bacteria in the blank-growth control group increased by 1.82 log_10_ CFU/mL. When the drug concentration was >1 MIC, the antibacterial effect showed different degrees of inhibition. The bacteria at 2, 4, 8, and 16 MIC decreased by 0.742, 0.858, 2.03, and 2.65 log_10_ CFU/mL, respectively. At 32 MIC, the amount of bacteria decreased by 3.16 log_10_ CFU/mL, thereby achieving a bactericidal effect. In summary, the antibacterial effect of tilmicosin against *M. hyopneumoniae* was more obvious with an increase in the drug concentration.

**Figure 2.**
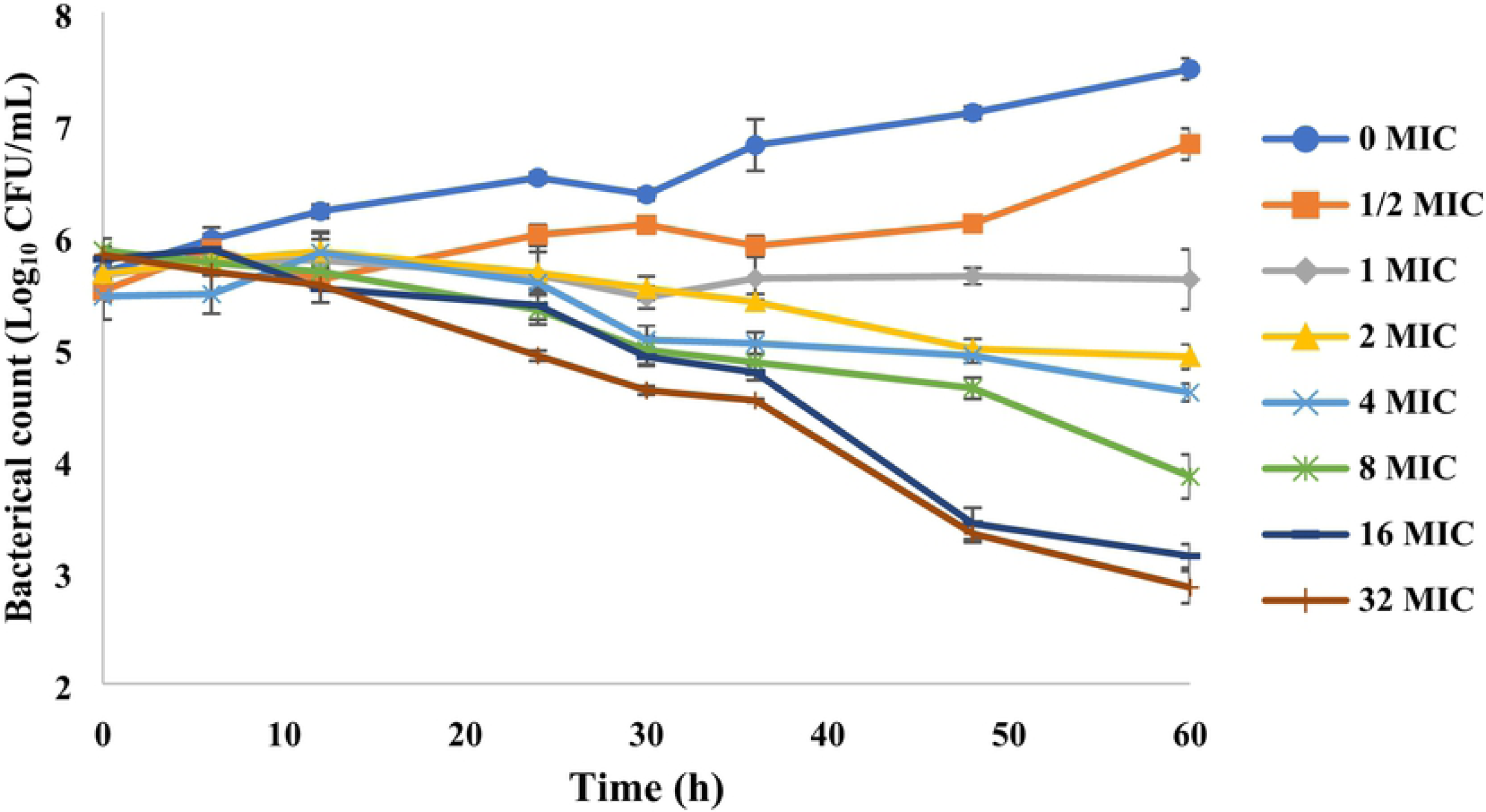
Time–kill studies of tilmicosin against *M. hyopneumoniae* at constant concentrations. MIC, minimum inhibitory concentration; CFU, colony-forming units. Data points represent the geometric mean values of three experiments.

In the *in vitro* dynamic model, the bactericidal curve of tilmicosin at different clinically recommended doses is shown in **Figure 3**. Within 0 h to 36 h, the number of bacteria under different drug doses did not decrease significantly, which suggested that the antibacterial effect of tilmicosin against *M. hyopneumoniae* took a long time. With an increase in the drug dose, tilmicosin showed increased activity against *M. hyopneumoniae*. The *M. hyopneumoniae* count was reduced from 3.61 to 5.11 log_10_ CFU/mL upon incubation for 96 h in a dynamic model with a dose of 40–200 mg, and achieved bactericidal activity.

**Figure 3.**
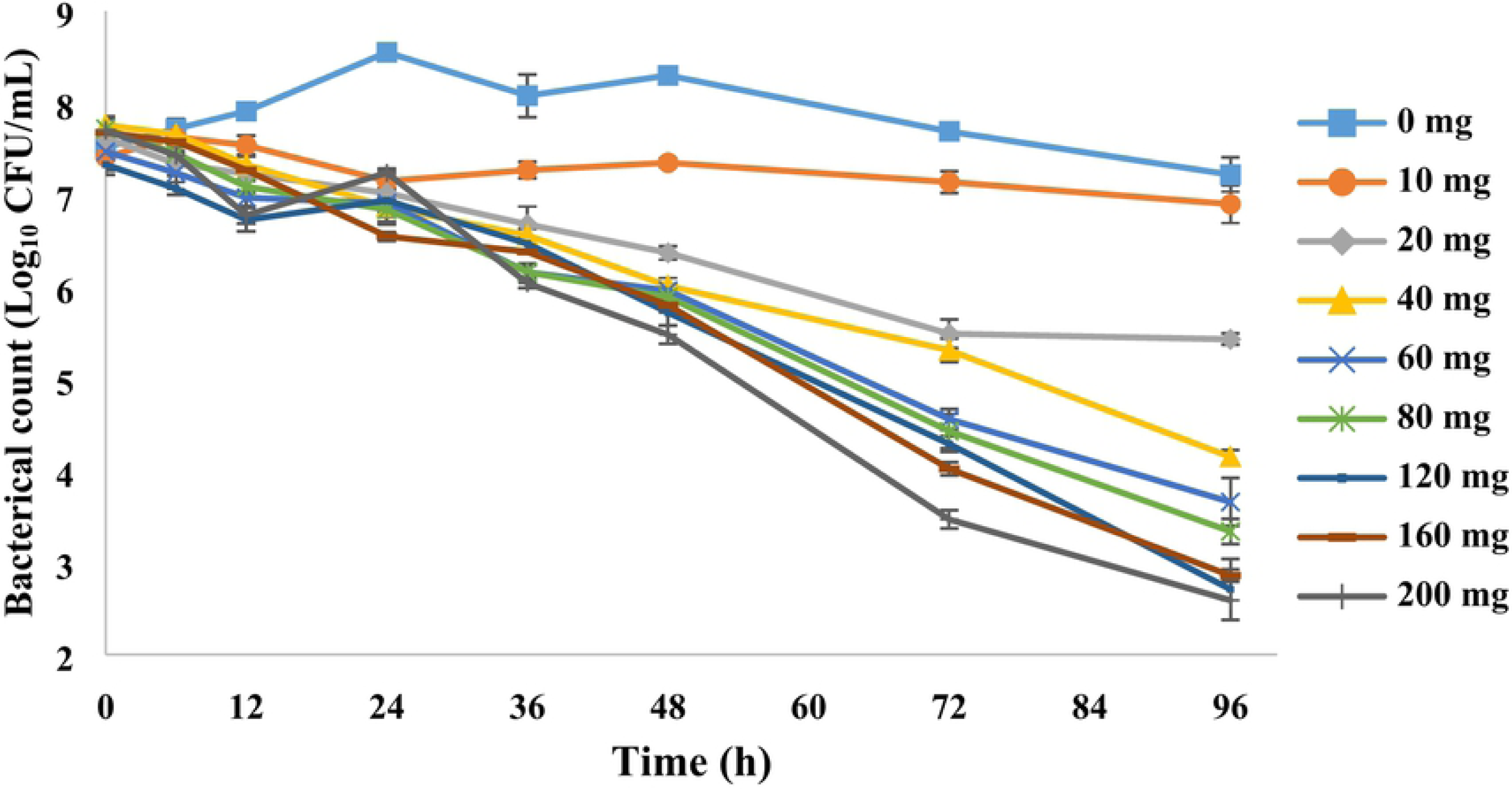
Dynamic time–kill curves were plotted at eight doses of tilmicosin. Data points represent the geometric mean values of three experiments.

### 3.3 PK in the *in vitro* dynamic model

The time–concentration curves for the *in vitro* dynamic model are shown as **Figure 4**. The PK parameters are summarized in **Table 1**. A standard curve was constructed by addition of a specific concentration of tilmicosin to the drug-free medium at concentrations ranging from 0.005 μg/mL to 0.5 μg/mL (R^2^ > 0.99). The relative error of elimination half-life and absorption half-life was −5.6 and 9.28%, respectively, both of which were within the normal range of ±15%. The LoD and LoQ for the developed method was 0.5 ng/mL and 1 ng/mL, respectively.

**Table 1.**
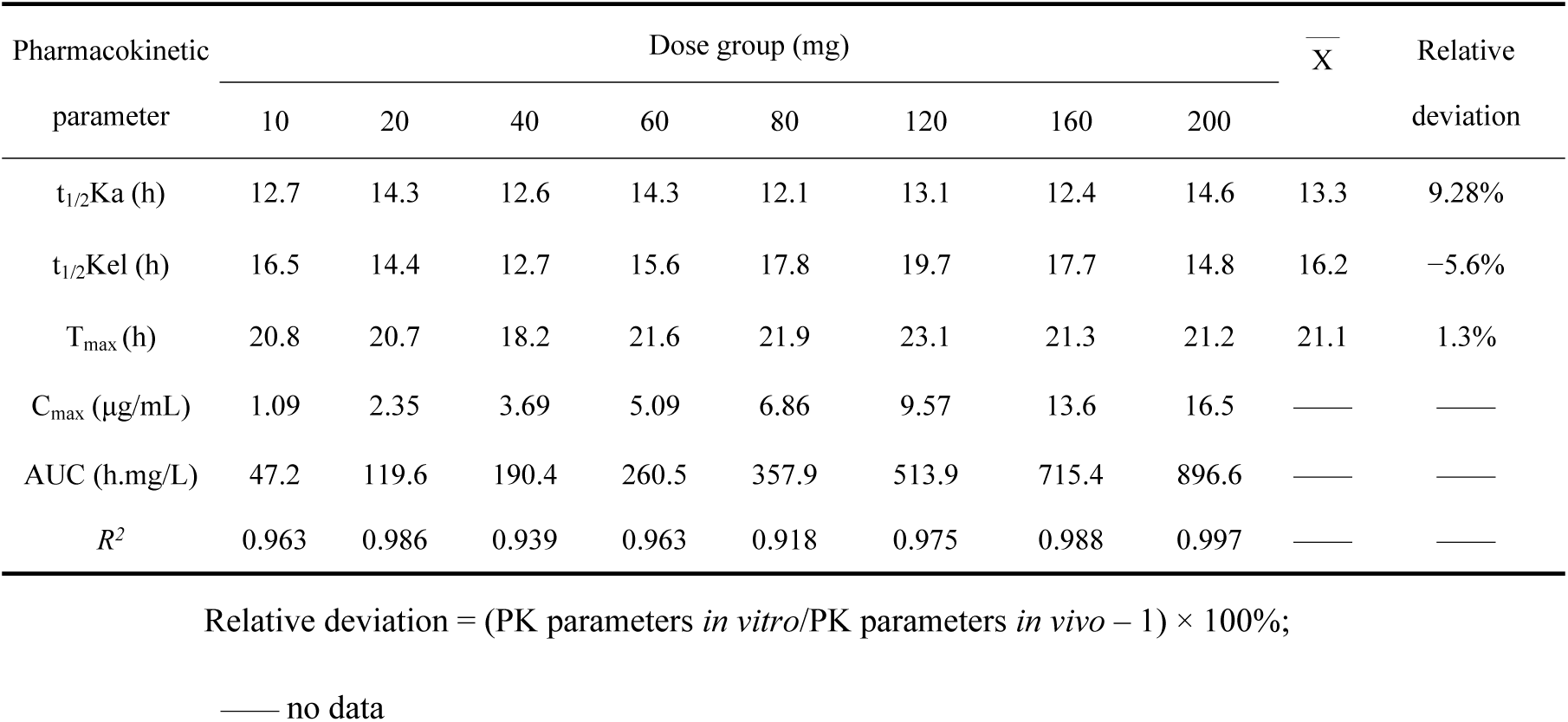
Main pharmacokinetic parameters of different tilmicosin doses in the *in vitro* PK/PD model.

**Figure 4.**
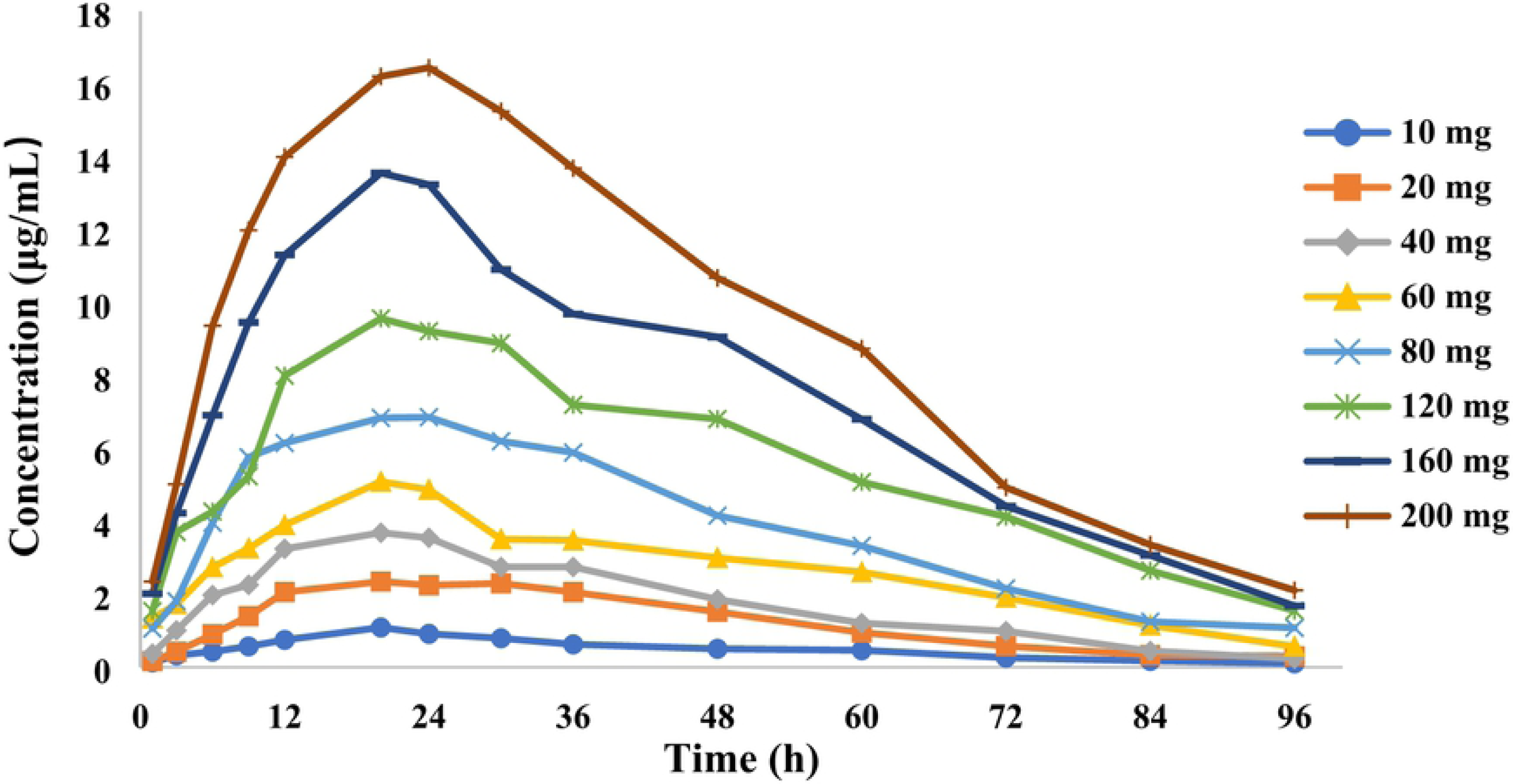
Concentration–time curves of eight doses of tilmicosin according to simulation of the timicosin concentration in the fluid lining the lung epithelia of pigs in the *in vitro* dynamic model.

### 3.4 Modeling and analyses of PK/PD

The relationships between antibacterial effect and PK/PD parameters of our dynamic model are shown in **Figure 5**. According to simulation of the inhibitory sigmoid *E*_*max*_ model, the correlation coefficient between AUC_0−96 h_/MIC, C_max_/MIC and the antibacterial effect was both 0.99. C_max_/MIC and AUC_0–96 h_/MIC were the best-fit PK/PD parameters for predicting the antibacterial activity of tilmicosin against *M. hyopneumoniae*, which suggested that tilmicosin had concentration-dependent activity. The estimated value for C_max_/MIC and AUC_0–96 h_/MIC for 2log_10_ (CFU/mL) reduction and 3log_10_ (CFU/mL) reduction from baseline was 1.44 and 1.91, and 70.55 h and 96.72 h, respectively, during a 96-h treatment period of tilmicosin. The obtained parameters of *E*_0_, *E*_max_, and *EC*_*50*_, and the Hill coefficient are listed in **Table 2**.

**Table 2.** Estimation of PK/PD parameters (data are derived from the *E*_max_ model).

| PK/PD parameter | $E_{max}$ | $EC_{50}$ | $E_0$ | Hill's | $R^2$ |
| --- | --- | --- | --- | --- | --- |
|  | (log<sub>10</sub> CFU/mL) |  | (log<sub>10</sub> CFU/mL) | slope |  |
| AUC<sub>0-96 h</sub>/MIC (h) | -0.30 | 91.93 | -5.28 | 1.65 | 0.99 |
| $C_{max}$/MIC | -0.30 | 1.80 | -5.17 | 1.89 | 0.99 |

**Figure 5.**
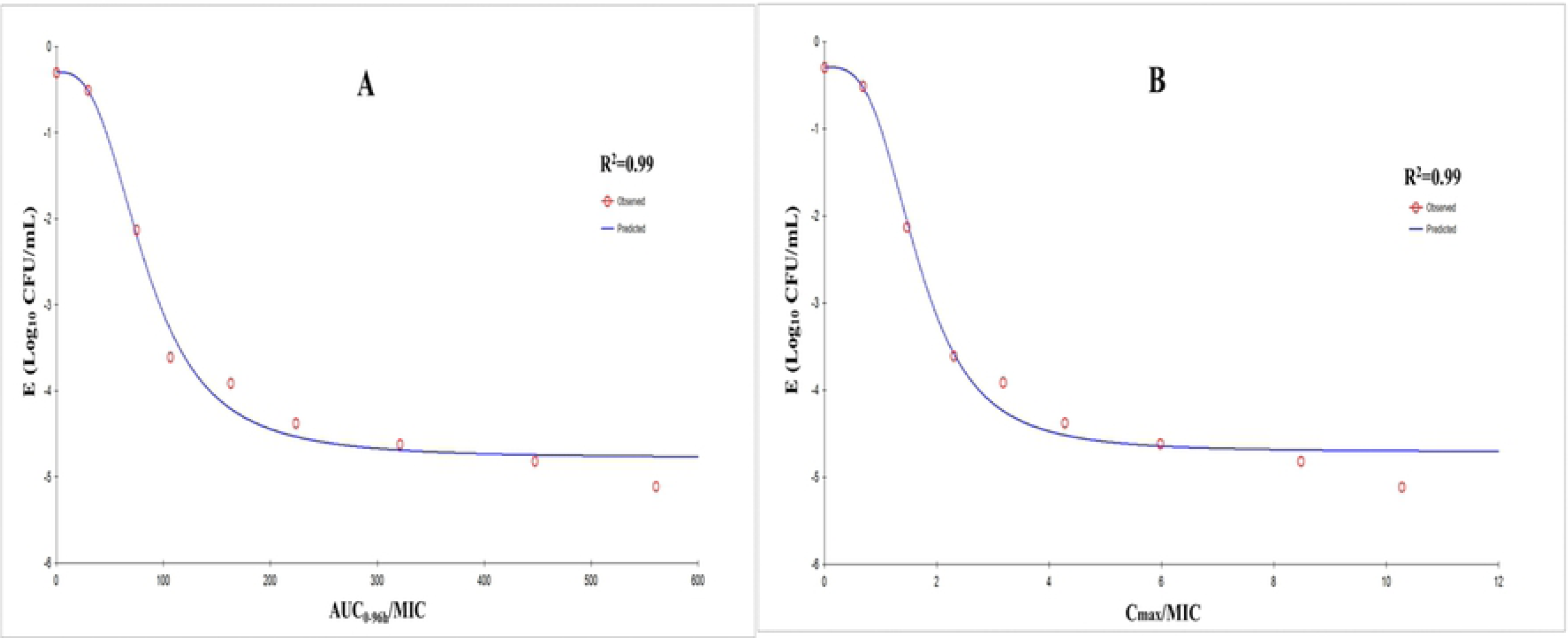
*E*_max_ relationships for three PK–PD parameters *versus* the antimycoplasmal effect using curves. **A**. AUC_0–96h_/MIC–antimycoplasmal effect; **B**. C_max_/MIC–antimycoplasmal effect; *R*^2^ is the correlation coefficient; *E*_max_ is the sigmoid maximum effect; PK, pharmacokinetic; PD, pharmacodynamic; AUC, area under the concentration–time curve; MIC, minimum inhibitory concentration; C_max_/MIC is the peak concentration by MIC.

### 3.5 Susceptibility testing and mutation analyses

Four *M. hyopneumoniae* strains (M1, M2, M3, and M4) with reduced sensitivity to tilmicosin were isolated from the four dose groups (10, 60, 80, and 120 mg). The MIC of these strains to tilmicosin ranged from 25.6 μg/mL to 1638.4 μg/mL. Moreover, the MIC of strains M3 and M4 was significantly higher than the MIC of standard strains. **Table 3** shows the changes in sensitivity of these strains to eight antimicrobial agents. The susceptibility of these strains to tylosin, erythromycin and lincomycin was also reduced significantly. For sequencing analyses of 23S rRNA (**Table 4**), we measured the gene sequences of regions V, L4, and L22, and compared them with standard strains. An acquired A2058G transition in region V was found only in the resistant *M. hyopneumoniae* strains (M3, M4) and a resistance mutation was not found in L4 or L22.

**Table 3.** MIC of six antimicrobial agents against *M. hyopneumoniae* (Mhp) and M1–M4 strains.

| Strain | MIC value of antibiotics (µg/mL) |  |  |  |  |  |  |  |
| --- | --- | --- | --- | --- | --- | --- | --- | --- |
|  | tilmicosin | tylosin | erythromycin | tiamulin | doxycyclin | enrofloxacin | amikacin | lincomycin |
| Mhp | 1.6 | 0.0625 | 0.64 | 0.08 | 0.0625 | 0.025 | 1.6 | 0.125 |
| M1 | 25.6 | 0.125 | >40.96 | 0.08 | 0.0625 | 0.025 | 1.6 | 0.125 |
| M2 | 819.2 | 8 | >40.96 | 0.08 | 0.0625 | 0.025 | 1.6 | 64 |
| M3 | 1638.4 | 16 | >40.96 | 0.08 | 0.0625 | 0.025 | 1.6 | 256 |
| M4 | 1638.4 | 16 | >40.96 | 0.16 | 0.0625 | 0.025 | 1.6 | 256 |

M1 (10 mg), M2 (60 mg), M3 (80 mg) and M4 (120 mg) strains were selected from seven doses, respectively.

**Table 4.**
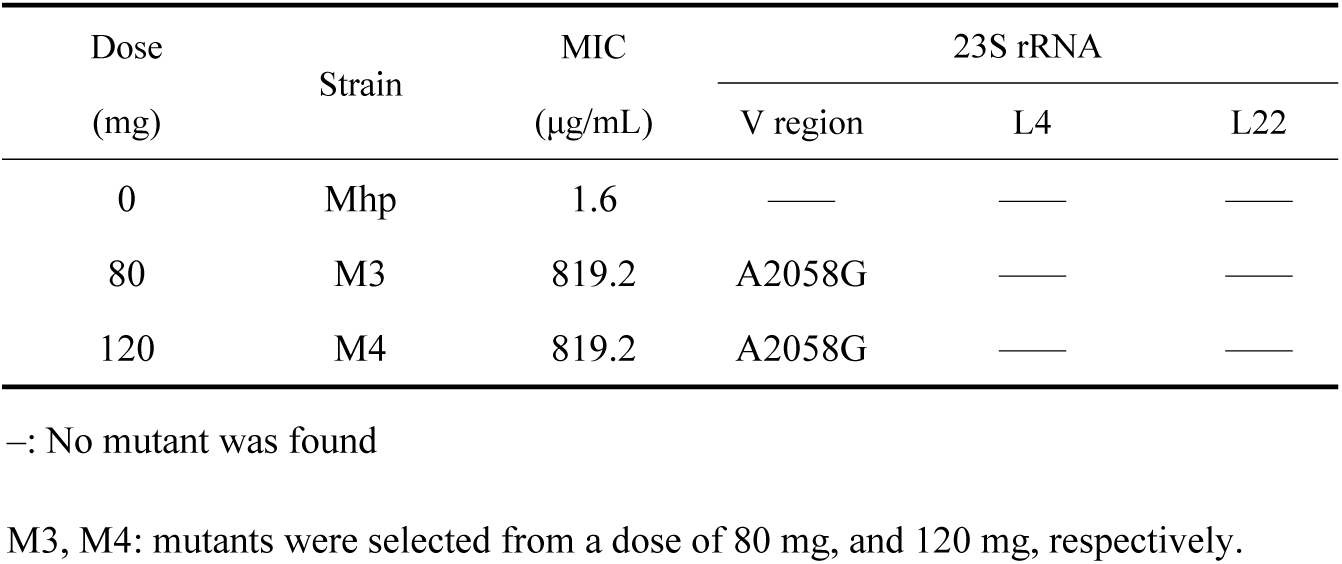
Tilmicosin susceptibility and identification of resistant mutants associated with different doses of tilmicosin.

| Dose<br>(mg) | Strain | MIC<br>(µg/mL) | 23S rRNA |  |  |
| --- | --- | --- | --- | --- | --- |
|  |  |  | V region | L4 | L22 |
| 0 | Mhp | 1.6 | — | — | — |
| 80 | M3 | 819.2 | A2058G | — | — |
| 120 | M4 | 819.2 | A2058G | — | — |

—: No mutant was found
M3, M4: mutants were selected from a dose of 80 mg, and 120 mg, respectively.

## 4. Discussion

Mycoplasmal pneumonia caused by *M. hyopneumoniae* infection is a major problem in the worldwide pig industry. If *M. hyopneumoniae* is mixed with other pathogenic microorganisms, porcine respiratory disease complex can occur [1, 21].

Rational use of antibiotics is the main measure to prevent and treat mycoplasmal pneumonia. However, *M. hyopneumoniae* presents technical difficulties in culturing and agar-plate enumeration both in the culture environment and in the addition of nutrients [22]. *M. hyopneumoniae* grows slowly and is inhibited competitively by other bacteria, so isolation of *M. hyopneumoniae* strains is difficult [23]. Hence, studies on the PK/PD interaction of tilmicosin against *M. hyopneumoniae* are scarce.

The *in vitro* dynamic model has been used widely to study PK/PD interactions. Tam et al. [24] investigated the PK/PD relationship of polymyxin B against *Pseudomonas aeruginosa* using an *in vitro* dynamic model. They showed that polymyxin B had rapid and concentration-dependent bactericidal activity against *P. aeruginosa*, and that the effect was diminished if the amount of bacteria was increased. Also, study of the effect of doxycycline against *Mycoplasma gallisepticum* in an *in vitro* model led to determination of the optimal PK/PD parameters to prevent drug resistance [25].

Here, this study reported on an *in vitro* PK/PD model of *M. hyopneumoniae* which simulated the PK of tilmicosin in bronchoalveolar lavage fluid after a single oral dose. The advantage of this model is that, when there is great difficulty in establishing an animal-infection model, the interaction between the drug and bacteria can be elucidated, and the change in drug sensitivity of the pathogen can be observed to study the drug-resistance mechanism.

Vicca et al. [26] determined the *in vitro* susceptibility of *M. hyopneumoniae* field isolates by a broth microdilution method. They showed that the MIC range of tilmicosin against *M. hyopneumoniae* was 0.25–16 μg/mL. Felde and colleagues reported that the MIC of tilmicosin against *M. hyopneumoniae* was 0.25–64 μg/mL [27]. Compared with those two studies, the MIC determined in the present study was within a reasonable range. The recommended turbidity of MIC testing against veterinary *Mycoplasma* species is 10^3^–10^5^ CFU/mL [28]. However, the turbidity of MIC testing used in our experiments was 10^5^–10^7^ CFU/mL and the inoculum used in the *in vitro* dynamic model experiment was high (10^7^ CFU/mL).

There were three main reasons for choosing a high inoculum. First, turbidity has little effect on the growth of *Mycoplasma* species or MIC determination [29]. Second, tilmicosin accumulates mainly in the lungs, so the concentration in lung tissues is much higher than that in plasma. If the drug concentration in the lungs is simulated, use of low bacterial counts is eliminated rapidly. Third, mutant subpopulations are present at low frequencies (10^−6^ to 10^−8^) [30]. Therefore, a high inoculum may increase the likelihood of monitoring mutant strains and resistance mechanisms. The binding rates of plasma proteins were not considered in the *in vitro* dynamic assay, but also because the tilmicosin concentration in plasma was much lower than that in the lungs. Using the drug concentration in plasma as a reference is not reasonable.

The time–kill curves in **Figure 2** showed that the antibacterial effect was more obvious when increasing the drug concentration (1–32 × MIC). The same situation also appeared in the *in vitro* dynamic time–kill curve (**Figure 3**). When the tilmicosin dose reached 40 mg, the bacteria decreased by 3.6 log_10_ CFU/mL. We noted a slight decrease in the amount of bacteria in the blank-growth control group and low-dose group (10 mg). There are two reasons for this result. First, the nutritional conditions required for *M. hyopneumoniae* growth were reduced slightly due to the long period of the dynamic model test. Second, in the low-dose group, there was a post-antibiotic sub-MIC effect (PA-SME), and the amount of bacteria may decrease slightly under a PA-SME.

The results of *E*_*max*_ model-fitting confirmed that the effect of tilmicosin on *M. hyopneumoniae* was concentration-dependent, and the parameters AUC_96h_/MIC and C_max_/MIC had high correlation with antibacterial effects (*R*^*2*^ = 0.99). Most studies have shown that the %T > MIC parameter of macrolides is associated significantly with antimicrobial activity [31, 32]. However, the antibacterial activity of azithromycin with a long elimination half-life is related to the AUC_24 h_/MIC parameter [33]. Therefore, the antibacterial activity of antibiotics is not static: it is dependent upon the characteristics of drugs and bacteria. The same antibiotic has different types of action on different bacteria [34]. In This study, it showed that the antibacterial effect of gentamicin on *Staphylococcus aureus* was time-dependent, while the antibacterial effect on *Pseudomonas aeruginosa* was concentration-dependent.

This study screened resistant strains of *M. hyopneumoniae* using drug-containing agar plates. And obtained four strains with significantly reduced sensitivity to tilmicosin. Studies on the mechanism of resistance of *Mycoplasma* species to macrolides are limited to mutations in drug-target molecules and efflux of antibacterial-active substances. We found that the A2058G mutation in domain V of the 23S rRNA gene (M3 and M4) was associated with resistance. No resistance-related differences were found in the ribosomal proteins L4 and L22. Studies have also shown that macrolide-resistant *M. hyopneumoniae* strains isolated from animals have mutations at positions 2057, 2058, 2059, or 2064 in the 23S rRNA gene [9, 10].

This study had two main limitations. First, all experiments were undertaken *in vitro*. Although the effects of the drug on bacteria were studied carefully and thoroughly, the effects of the immune system of an animal on microorganisms were not considered. Second, only one standard strain of *M. hyopneumoniae* was tested, and testing of clinical isolates is necessary to confirm our findings.

## 5. Conclusions

This was the first study on the PK/PD relationship of tilmicosin against *M. hyopneumoniae*. In the *in vitro* dynamic model, tilmicosin produced a maximal anti-*M. hyopneumoniae* effect of a 5.11 log_10_ (CFU/mL) reduction. The antibacterial effect of tilmicosin was concentration-dependent, and the best-fit PK/PD parameters were the AUC_0–96 h_/MIC and C_max_/MIC (R^2^ = 0.99). The estimated value for C_max_/MIC and AUC_0–96 h_/MIC for 2log_10_ (CFU/mL) reduction and 3log_10_ (CFU/mL) reduction from baseline was 1.44 and 1.91, and 70.55 h and 96.72 h, respectively. The A2058G mutation in region V of the 23S rRNA gene was found in M3 and M4 strains. These results provide a reliable reference for animal experiments *in vivo*, and may help in the design of more rational treatment for *M. hyopneumoniae* infection.

## Acknowledgments

This work was supported by the National Key Research and Development Program of China (2016YFD0501300 and 2016YFD0501310).

## Author contributions

Methodology, software use, validation, analyses, data curation, manuscript preparation, manuscript reviewing/editing, visualization, and project administration were done by ZH. ZH, HZ, ZH and XX contributed to the investigation. Resources were provided by XG, XS, HY, and HD. Supervision was provided by HD, who also acquired the funding.

